# The CFII components PCF11 and Cbc change subnuclear localization as cells differentiate in the male germ line adult stem cell lineage

**DOI:** 10.1101/2025.07.22.666077

**Authors:** Iliana Nava, Margaret T. Fuller, Lorenzo Gallicchio

## Abstract

Stage specific increased expression of PCF11 and Cbc, the two components of Cleavage Factor complex II, contribute to developmentally regulated 3’UTR shortening due to alternate polyadenylation of nascent RNA molecules in *Drosophila* spermatocytes compared to spermatogonia. Here, we show that both Cbc and PCF11 change subnuclear localization during male germ cell differentiation, from homogeneous in the nucleus in spermatogonia and early spermatocytes to concentrated around the nucleolus in later spermatocyte stages.

## Description

Co-transcriptional Cleavage and Polyadenylation of nascent RNA molecules is the final step of transcription and is required for the newly transcribed mRNAs to be exported to the cytoplasm (Rodríguez-Molina et al., 2023). A multi subunit cleavage machinery complex responsible for making the 3’end cut directs cleavage and polyadenylation to specific sites on the nascent RNA by recognizing specific hexametric RNA motifs termed Polyadenylation Signals (PAS). The canonical PAS (AAUAAA) and some variants considered “strong” are more frequently processed by the cleavage machinery than “weak” variants or non-canonical PAS (Gallicchio et al., 2023; Gruber & Zavolan, 2019; Neve et al., 2017). Genes in multicellular eukaryotes usually possess multiple PAS motifs, and the same genetic locus often gives rise to alternative mRNA isoforms that differ at their 3’ end due to a different choice of which PAS is utilized, a process named Alternative Polyadenylation (APA). An alternate PAS located upstream of a stop codon can result in a truncated or variant protein. Most commonly, alternate PAS sites located in the 3’ untranslated region (3’UTR) can result in mRNA isoforms that differ in 3’UTR length and content. As 3’UTRs are a hub of cis-regulatory sequences that attract *trans*-acting factors such as microRNAs and RNA binding proteins, changes in the 3’UTR can affect mRNA stability, localization and translation (Gallicchio et al., 2023). Thus, APA can profoundly affect what proteins are expressed by simply changing the site at which transcripts are terminated.

Changes in APA have been implied in numerous disease states (Davis et al., 2022; Gruber & Zavolan, 2019; Kurozumi & Lupold, 2021; Ren et al., 2020; Tian & Manley, 2016; Venkat et al., 2020; Y. Zhang et al., 2021). For example, certain cancer cell lines tend to express mRNAs with shorter 3’UTRs, causing loss of repressors (i.e. miRNAs) binding sites and subsequent abnormal upregulation of the encoded proteins(Mayr & Bartel, 2009; Y. Zhang et al., 2021). Recently, APA has been shown to be a feature of many healthy biological processes, and that APA must be tightly regulated to maintain cell and tissue homeostasis (Gallicchio et al., 2023; Kasowitz et al., 2018; Smibert et al., 2012; Ulitsky et al., 2012; Vallejos Baier et al., 2017; Yang et al., 2022).

Developmentally regulated processing of alternate PAS sites can program cell type specific changes in 3’UTRs that profoundly alter where and when certain proteins are expressed. For example, differentiating neurons tend to express mRNA isoforms with longer 3’UTRs due to APA (Oktaba et al., 2015; Vallejos Baier et al., 2017). This may be particularly important for proper sub-cellular localization of specific mRNAs in neurons (Arora et al., 2022). Conversely, differentiating spermatocytes tend to express mRNA isoforms with shorter 3’UTRs compared to the transcripts expressed from the same genes in spermatogonia (Berry et al., 2022; Li et al., 2016; Shan et al., 2017). In *Drosophila* spermatocytes, more than 500 genes switch from a distal cleavage site (used in spermatogonia) to a more proximal one due to developmentally regulated APA (Berry et al., 2022) (Fig 1 A-B). Strikingly, the resulting switch in 3’UTR length correlated with changes in protein expression: in some cases, the protein was not detected in spermatogonia but highly translated in spermatocytes, while in others the encoded protein was translated in spermatogonia but not detected in spermatocytes (Berry et al., 2022).

**Fig 1.**
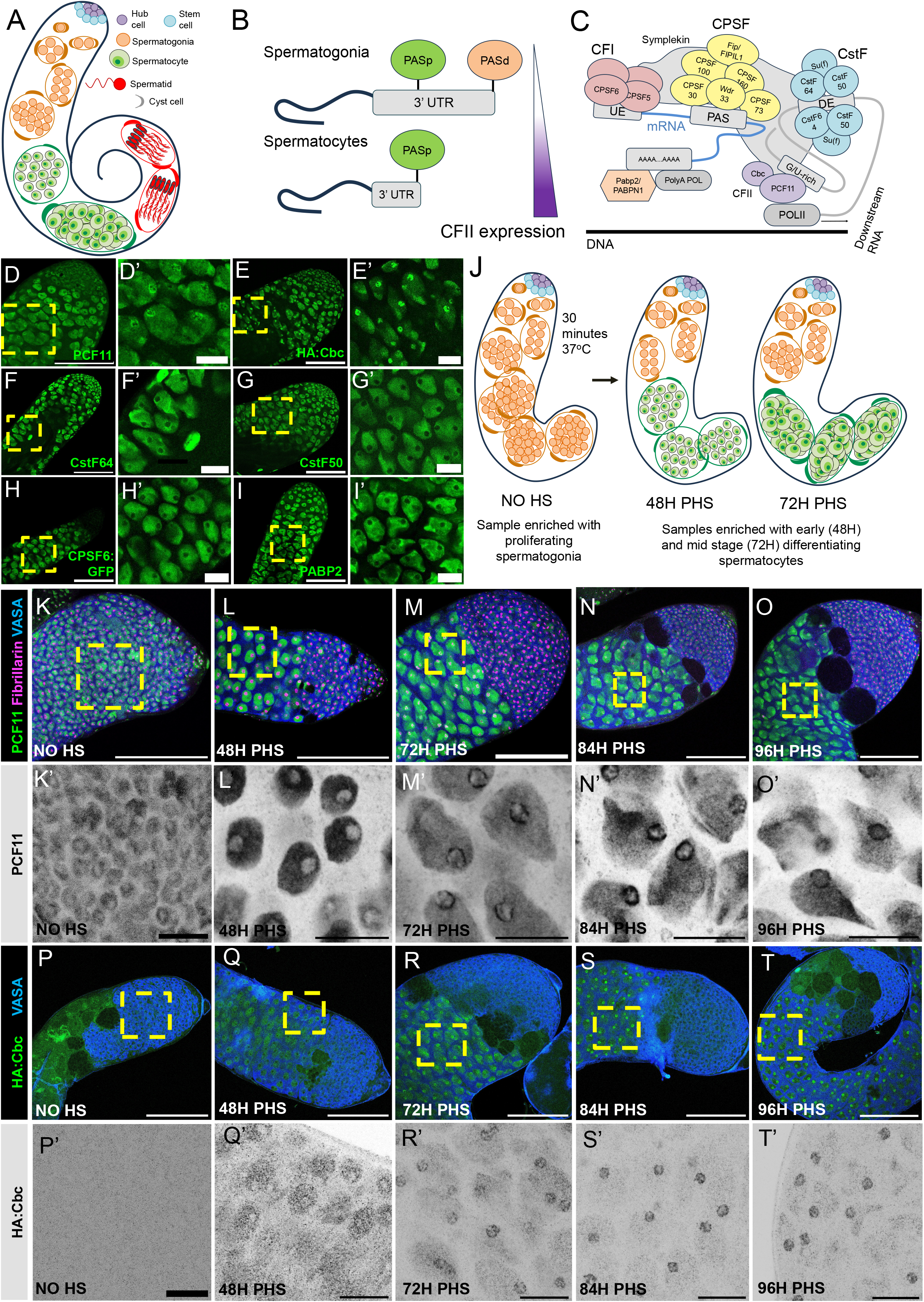
PCF11 and Cbc change localization from homogeneous in the nucleoplasm to more concentrated at the nucleolar periphery as spermatocytes grow. (A) Schematic of a *Drosophila* testis. (B) Diagram of the APA switch observed as spermatogonia differentiate into spermatocytes. PASd (orange): the distal PAS processed in spermatogonia. PASp (green): the proximal PAS processed in spermatocytes. The APA switch is at least partially regulated by increased expression of CFII in spermatocytes compared to spermatogonia. (C) Diagram of the cleavage machinery and its complexes: CPSF (yellow), CstF in (blue), CFI (red) and CFII in (purple). UE = Upstream Element, DE = Downstream Element. (D-I) Apical tips of fixed *Drosophila melanogaster* testes immune-stained for: (D-D’) PCF11, (E-E’) HA:Cbc, (F-F’) CstF64, (G-G’) CstF50, (H-H’) CPSF6:GFP, (I-I’) PABP2. Scalebars are 100 uM (D-I) and 20 uM (D’-I’). Genotypes: *w1118* (D-F-G-I), *HA:Cbc* (E), *CPSF6:GFP* (VDRC 318105) (J) Diagram of the heat shock time course system for synchronous spermatocyte differentiation. PHS = post heat shock. (K-O) Apical tips of fixed *Drosophila melanogaster* testes immuno-stained with antibodies against: PCF11 (green), Fibrillarin (magenta) and VASA (blue). Timepoints: (K) no heat shock, (L) 48H PHS, (M) 72H PHS, (N) 84H PHS, (O) 96H PHS. Scalebars are: 100 uM (K-O) and 20 uM (K’-O’). (P-T) Apical tips of fixed *Drosophila melanogaster* testes immune stained with antibodies against: HA:Cbc (green), and VASA (blue). Timepoints: (P) no heat shock, (Q) 48H PHS, (R) 72H PHS, (S) 84H PHS, (T) 96H PHS. Scalebars are: 100 uM (P-T) and 20 uM (P’-T’).

The tendency to cleave nascent transcripts at more proximal PASs in *Drosophila* spermatocytes was in part due to cell type specific upregulation of specific components of the cleavage machinery. The cleavage machinery is composed of 4 major protein complexes: CPSF (Cleavage and Polyadenylation Specificity Factor), CstF (Cleavage Stimulation Factor), CFI (Cleavage factor I) and CFII (Cleavage Factor II), plus the non-complex proteins Symplekin, PolyA Polymerase and PABP2 (Gallicchio et al., 2023). Genetic experiments showed that high levels of the two components of CFII, PCF11 and Cbc, were required in spermatocytes to direct cleavage at a proximal site rather than the PAS used in spermatogonia for over 200 genes (Gallicchio et al., 2024). Cbc had been recently reported to change sub-nuclear localization as spermatocytes differentiate in *Drosophila* (Wu et al., 2021). While in spermatogonia Cbc was homogenously distributed in the nucleus, in late spermatocytes Cbc protein concentrated particularly in and around the nucleolus (Wu et al., 2021) (Fig. 1 D).

Here we show that PCF11 also changes sub-nuclear localization as germ cells differentiate (Fig. 1 E). We did not observe such re-colocalization to the nucleolus for several other cleavage factors tested: CstF50, CstF64, PABP2 and CPSF6 (Fig. 1 F-G-H-I). Analysis of *Drosophila* testes enriched for specific germ cell stages through an *in vivo* synchronous differentiation strategy identified the stage of spermatocyte differentiation at which the switch of sub-nuclear localization of Cbc and PCF11 occurred. During spermatogenesis, proliferating spermatogonia undergo four rounds of transient amplifying division, producing a clone of interconnected germ cells enclosed in an envelope of two squamous epithelial somatic support cells, forming a cyst (Fig. 1 A). The 16 cells then enter premeiotic S phase and initiate the spermatocyte program of cell growth and transcriptional activation. The switch from spermatogonial proliferation to spermatocyte differentiation relies on function of the *bag of marbles (bam*) gene, with Bam protein required to reach a specific threshold to trigger the switch (Insco et al., 2009). Flies that completely lack *bam* have testes filled with cysts containing spermatogonia that undergo several extra rounds of mitotic division then die (Harris et al., 2025; Insco et al., 2009). Providing a brief pulse of Bam expression under control of a heat shock inducible promoter can induce *bam*^-/-^ spermatogonia to differentiate into spermatocytes, then eventually spermatids and sperm (Jongmin Kim et al., 2017). Collecting samples at different time points after heat shock yields testes containing semi-synchronous cohorts of spermatocyte cysts at specific stages of maturation, depending on the time after heat shock (Fig. 1 J).

Immunofluorescence staining at different time points after heat shock revealed that as spermatocytes mature, PCF11 and Cbc, the two components of CFII, change sub-nuclear localization. In spermatogonia (no heat shock sample, Fig. 1 K-K’) PCF11 is homogenously distributed in the nucleoplasm. At 48H PHS testes had newly formed *bam*^-/-^ spermatogonia at the apical tip and the rest of the testes were filled with early, polar spermatocytes. At this stage, PCF11 and HA:Cbc were upregulated in spermatocytes compared to the juxtaposed spermatogonia but both proteins were homogeneously distributed in the spermatocyte nuclei and excluded from the nucleolus, similarly to spermatogonia (Fig. 1 L and Q). In testes fixed at 72 H PHS, both PCF11 and HA:Cbc showed higher immunofluorescence signal around the nucleolar periphery than in the general nucleoplasm (Fig. M and R). At later time points (84H and 96H PHS) the signal around the nucleolus intensified (Fig. 1 N, S, and O, T, respectively) (see SOM 1-4 for additional examples).

The nucleolus, a key membrane-less nuclear compartment, is the site of processing of ribosomal RNA (rRNA) and ribosome assembly. Recently, transcription of rDNA spacer regions by PolII has been shown to have an active role in rRNA production and nucleolar architecture (Abraham et al., 2020; Caudron-Herger et al., 2016; Khosraviani et al., 2024). Consistent with active PolII dependent transcription near the nucleolus, the tTAFs (testis specific TATA binding protein Associated Factors), required for robust transcription at spermatocytes specific promoters, have been shown to concentrate at the nucleolus in spermatocytes (Chen et al., 2005). As differentiating spermatocytes grow 25 times in volume and also need to assemble many ribosomes to endow sufficient capability to translate the many new mRNAs necessary for the post meiotic morphological transformation to mature sperm, it is likely that transcription of rRNAs and their PolII dependent spacer regions is strongly upregulated in spermatocytes. PCF11 has been shown to bind to the C terminal domain of PolII and to regulate transcription termination (Kamieniarz-Gdula et al., 2019; Sadowski et al., 2003; Z. Zhang & Gilmour, 2006). We propose that the concentration of PCF11 and Cbc around the perimeter of the nucleolus in growing spermatocytes may be due to binding pf PCF11 to active PolII, bringing Cbc with it. Whether PCF11 and/or Cbc have an active role in regulating rRNAs processing or transcription termination of rDNA spacer regions remains to be elucidated.

## Methods

### Drosophila lines and husbandry

Flies were raised at either room temperature or 25°C except as described for the heat shock time course. For HS time course experiments, *bam*^*1*/Δ86^ mutants carrying a *bam* transgene under the control of a heat shock promoter were heat-shocked at 37°C for 30 minutes in late larval and pupal stages, then returned to 25°C. Flies were collected at different time points after heat shock, and their testes were then dissected and fixed for immunofluorescence. Flies used in this study were: *w1118* (Bloomington *Drosophila* Stock Center [BL] 3605), *CPSF6:GFP* (Vienna *Drosophila* Resource Center [VDRC] 318105), *HA:cbc* (kindly gifted from the Wang lab (Wu et al., 2021), *hsBam/CyO;bam*^86^,*bamGal4/TM6B*(Jongmin Kim et al., 2017; D. McKearin & Ohlstein, 1995) and *bam*^1^/*TM6B* (D. M. McKearin & Spradling, 1990).

### Immunofluorescence and Imaging

Testes were dissected from 0-3 day old male flies in 1X PBS at room temperature in a cyclops dissecting dish for a maximum of 10 minutes, transferred to an Eppendorf tube and immediately fixed in 5% formaldehyde in 1X PBS for 30 minutes with rocking at room temperature. Testes were then rinsed twice with 1X PBS and permeabilized by rocking at room temperature for 30 minutes in 0.3% Triton X-100 and 0.6% sodium deoxycholate in 1× PBS. The testes were then washed 3 times 15 minutes each in PBST (1× PBS + 0.05% Tween-20) with rocking at room temperature then blocked in 1X Western Blocking Reagent (WBR, Roche 11921673001) in PBST for 30 minutes rocking at room temperature. After blocking testes were incubated in primary antibody in 1X WBR in PBST overnight at 4°C. The following day, testes were washed 3 times in PBST for 15 minutes each, rocking at room temperature, blocked again in 1X WBR in PBST for 30 minutes rocking at room temperature, and incubated for 2 hours with secondary antibody mix in 1X WBR in PBST with rocking at room temperature. Testes were then washed 3 times 15 minutes each in PBST, rocking at room temperature, then mounted on glass slides in 10 uL of ProLong Diamond antifade with DAPI (Thermo Fisher P36962). Antibodies used: rabbit anti-CstF64 (1:400; a kind gift from Zbigniew Dominski, (Skrajna et al., 2018)); guinea pig anti-CstF50 (1:300; a kind gift from John T. Lis, (Ni et al., 2008)); chicken anti-GFP (1:10,000; Abcam 13970); rabbit anti-PABP2 (1:500; a kind gift from Martine Simonelig (Benoit et al., 1999)); rabbit anti-PCF11 (1:200; a kind gift from David Scott Gilmour, (Z. Zhang & Gilmour, 2006)); rabbit anti-HA (1:500; Cell Signaling Technologies 3724), mouse anti-HA (1:200; Biolegends 901501); anti-Fibrillarin (1:300, Abcam ab4566); goat anti-Vasa (1:200; Santa Cruz Biotechnology sc-26877, discontinued); rat anti-Vasa (1:10 for supernatant or 1:100 for the concentrated version; both from Developmental Studies Hybridoma Bank antibody ID 760351). Secondary antibodies used were: donkey or goat anti-rabbit; anti-mouse; anti-guinea pig; anti-chicken; anti-rat; and anti-goat; all used at 1:500 and conjugated with either Alexa 488, Alexa 568, Alexa 594, or Alexa 647 (Thermo Fisher catalog). Slides were imaged on a Leica SP8 (Cell Science Imaging Facility (CSIF), Stanford University) and Images were processed with FIJI.

## Extended Figure Legend

**SOM1**. Apical tips of fixed *Drosophila melanogaster* testes collected 48H post heat shock (PHS) immuno-stained with antibodies against: PCF11 (green), and Fibrillarin (magenta). Scalebars are: 100 uM (A-B-C) and 20 uM (A’-B’-C’).

**SOM2**. Apical tips of fixed *Drosophila melanogaster* testes collected 72H post heat shock (PHS) immuno-stained with antibodies against: PCF11 (green), and Fibrillarin (magenta). Scalebars are: 100 uM (A-B-C-D) and 20 uM (A’-B’-C’-D’).

**SOM3**. Apical tips of fixed *Drosophila melanogaster* testes collected 84H post heat shock (PHS) immuno-stained with antibodies against: PCF11 (green), and Fibrillarin (magenta). Scalebars are: 100 uM (A-B-C-D) and 20 uM (A’-B’-C’-D’).

**SOM4**. Apical tips of fixed *Drosophila melanogaster* testes collected 96H post heat shock (PHS) immuno-stained with antibodies against: PCF11 (green), and Fibrillarin (magenta). Scalebars are: 100 uM (A-B-C) and 20 uM (A’-B’-C’).

## Acknowledgments

We thank all the members of the Fuller lab for sharing comments and suggestions on this work. We thank the Wang laboratory for the *HA:cbc* CRISPR tagged flies, the Lis laboratory for anti-CstF50, the Simonelig laboratory for anti-PABP2, the Dominski laboratory for anti-CstF64, and the Gilmour laboratory for anti-PCF11 antibodies. We thank the Cell Science Imaging Facility (CSIF) at Stanford University, particularly the CSIF manager Kitty Lee. The contents of this manuscript are solely the responsibility of the authors and do not necessarily represent the official views of the NCRR or the National Institutes of Health. We thank the Stanford Fly Media Center as well and all the staff working in the Department of Developmental Biology. This manuscript is dedicated to Rosa Ramirez, Ruben Nava and Ruben Nava Jr.

## Author contributions

I.N., L.G. and M.T.F. conceived the project. L.G. wrote and M.T.F. edited the manuscript, I.N. prepared the figure. L.G. designed and supervised the experimental work, I.N. performed experiments. Funding was acquired by L.G. and M.T.F.

