## Supplementary figures and images for "The CFII components PCF11 and Cbc change subnuclear localization as cells differentiate in the male germ line adult stem cell lineage"

### Supplementary Figures

#### **SOM1**. 48H PHS

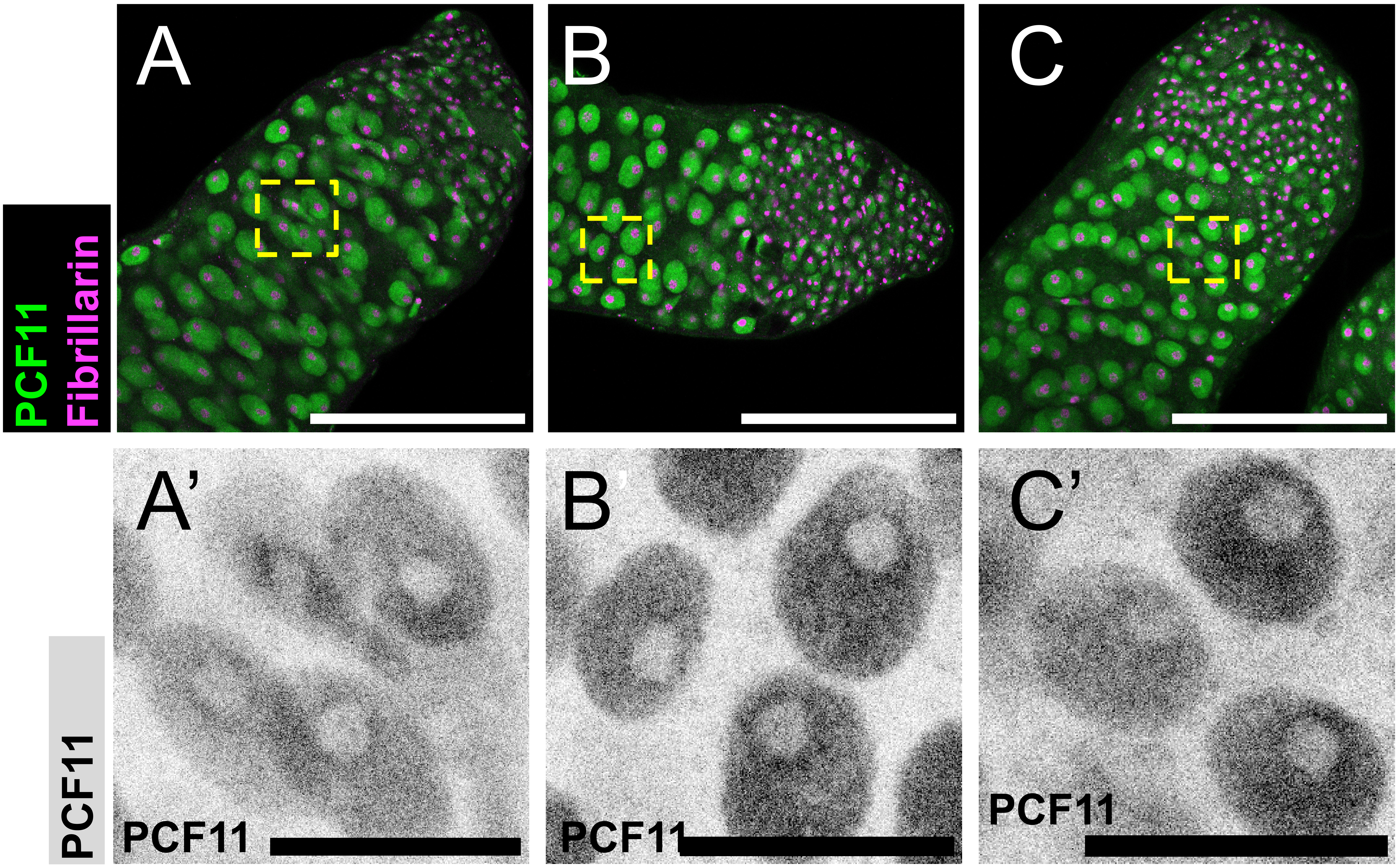

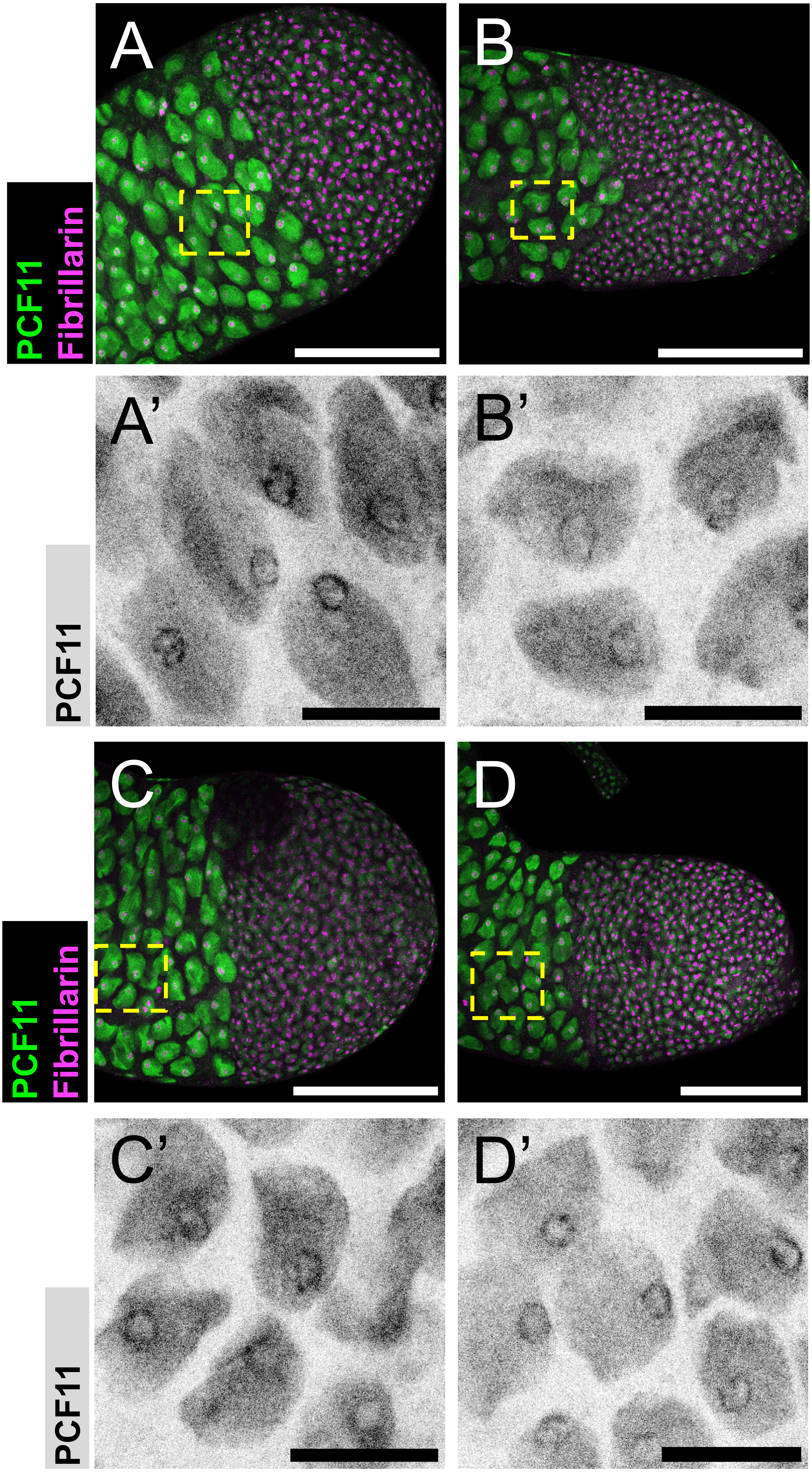

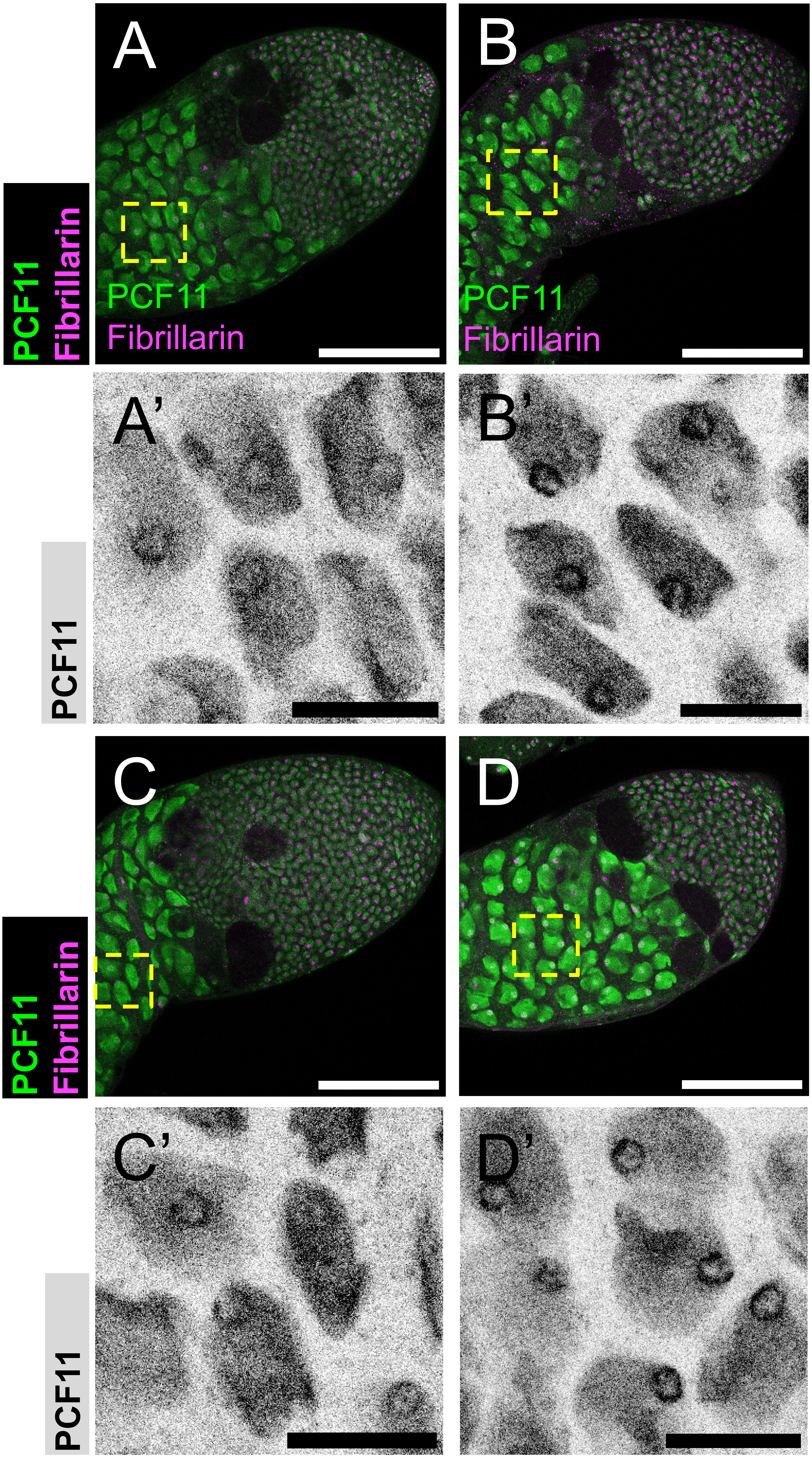

#### **SOM4**. 96H PHS

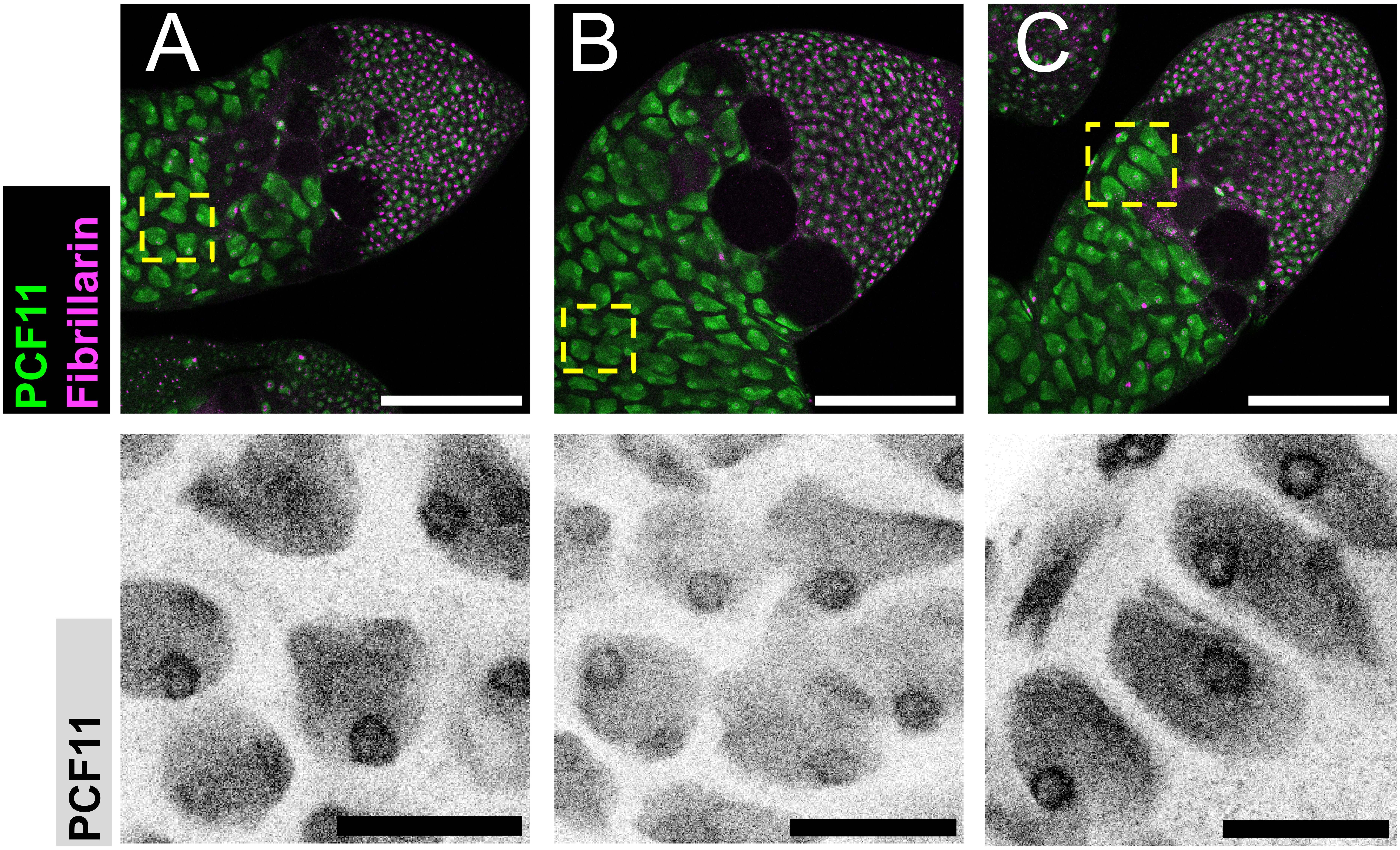
